## Supplemental Methods for "Population genomics of *Plasmodium ovale* species in sub-Saharan Africa"

**Supplemental Methods, Materials, and Figures**

**Sequencing data alignment, variant calling, and filtering**

Raw sequencing reads, including those of four isolates from *Joste et al.,* were trimmed of Illumina adapters using *Trimmomatic* v0.36^1^ before the quality of the reads was evaluated with *fastQC* v0.11.9*.**^2^* Paired reads from each isolate were aligned to their species’ corresponding dual reference genome using *bwa-mem2* v2.2.1^3^ before each alignment was sorted and given read group information using *picard* v2.26.11^4^ and deduplicated via *GATK* v4.4.0.0.^5^ After variant calling, variants within the following expanded gene families were masked via *vcftools**^6^* to reduce sequence error in repetitive genomic elements: *Plasmodium* interspersed repeat protein (*pir*), surfin-related subtelomeric protein 1 (*stp1*), early transcribed membrane protein (*etramp*), tryptophan-rich antigen (*tra*), *Plasmodium* exported protein (*phist)*, reticulocyte binding protein (*rbp*), ookinete surface protein (*osp*), 6-cysteine protein, KELT protein, and pm-fam-a protein.^7^ Tandem repeats across each species’ genome were identified and masked using Tandem Repeat Finder v4.09.1.^8^

**Complexity of infection (COI) calculation**

Variant call sets were limited to variants with a minor allele frequency greater than or equal to 5%. The McCOILR R package was employed to run THEREALMcCOIL on each sample set using 1000 markov chain monte carlo iterations, 100 burn-in iterations, a maximum COI of 25, a minimum number of sites for a sample to be included of 10, and a minimum number of samples for a site to be included of 10 (analysis was run on 13 and 16 *P. ovale curtisi* and *ovale wallikeri* isolates, respectively).^9^ This yielded 1000 estimates of complexity of infection, or the number of unique parasite clones found in each sample, and median COI per isolate was reported. COI distributions among *P. falciparum* isolates were calculated in the same manner. COI distributions among the three species were compared using a Kruskal-Wallis test, with Dunn’s multiple comparisons test employed for specific pairwise comparisons.^10^

To investigate the distribution of heterozygous SNPs in samples that were determined to be polyclonal, the variant set was filtered to sites with >10% minor allele frequency across the whole population and a within-sample minor allele frequency >50% for at least one sample. Then, for each polyclonal sample, remaining sites were filtered to those with a within-sample minor allele frequency greater than or equal to 5%.

**Principal components analysis**

After limiting to monoclonal samples (20 *Poc*, 23 *Pow*), variant call sets were filtered to remove sites with a minor allele frequency less than 5% using *vcftools* v0.1.15.^6^ Variants were processed using PLINK v1.90b6.21, first by pruning variants within windows of 50 variants between which the R^2^ value exceeded 0.3 (windows were shifted by steps of 5 variants for each pruning step).^11^ Then, principal components analysis was performed in PLINK, including extraction of the weights by variant. Data visualizations were performed in R using the *ggplot2* v3.4.4 package.^12^ ADMIXTURE v1.3.0 was employed to test for significant clustering among samples within each species.^13^

**Nucleotide diversity calculation**

For both ovale species and *P. falciparum*, the OrthoMCL database was searched for orthologous gene sets in which there was only one copy of each ortholog per genome.^14^ Sets were removed from the total pool of orthologous genes if they were partially or totally masked by the multigene family or extrachromosomal masks within an individual species, or if any orthologs did not have ≥5x coverage of the gene at least every 10 bases in at least 60% of isolates in each species. Among the remaining orthologs sets with sufficient coverage among all species, nucleotide diversity (𝛑) was calculated across the monoclonal samples of each species for each ortholog in the set using *vcftools*. The *P. falciparum* isolate set was composed of 19 monoclonal isolates selected to ensure one geographically-matched *Pf* sample for each *P. ovale* sample (**Supplemental Table 2**). Variant call sets in this analysis did not employ a minor allele frequency filter. Nucleotide diversity was compared between species among the remaining sets of orthologs using repeated measures ANOVA; pairwise comparisons were examined with two-tailed Tukey’s multiple comparisons test. To mitigate potential bias introduced by differing geographic coverage between the two ovale species (and by differences with the *P. falciparum* genome), nucleotide diversity was also calculated among a set of orthologue sets (identified among one-to-one *Poc*-*Pow* orthologues with strong coverage as detailed above) in geographically-matched *Poc* and *Pow* samples (n=11 each, **Supplemental Table 3**) and compared using Wilcoxon’s matched-pair signed rank test.^15^

**Identification of signatures of selection**

Variant call sets were limited to monoclonal isolates and sites which had no missingness in any samples. After employing the default minor allele frequency filter of >5%, *selscan* was used to calculate nS_L_ (a metric of directional selection) for all remaining variants in each species and the *norm* function used to normalize values in allele frequency bins.^16^ nS_L_ was chosen as it is more robust to the currently-unknown recombination rates across either *P. ovale* genome than metrics like iHS and because it can evaluate selection within a single population of organisms. Loci with an nS_L_ in the most extreme 0.5% were investigated in PlasmoDB for proximity to genes of interest within 10kb.^14^ Extended haplotype homozygosity (EHH) was calculated at select sites using *rehh.**^17^* Tajima’s D (a metric of possible directional and balancing selection) was calculated in 300 base pair sliding windows shifted by 10bp using all non-missing variants across all known genes in each genome using *vcf-kit.**^18,19^* Variants with positive Tajima’s D values in the top 0.5% of absolute values were investigated in PlasmoDB.

**Supplemental Figures and Tables**

| **Sample ID** | ***po18S* Ct value** | ***P. ovale* Species** | ***P. ovale* species Ct value (*Poc, Pow)*** | **Technique** | **Country of Origin** | **Study Site** | ***Pf* Coinfection** | **1x Cov. (%)** | **5x Cov. (%)** | **10x Cov. (%)** | **Average Mean Depth** |
| --- | --- | --- | --- | --- | --- | --- | --- | --- | --- | --- | --- |
| **DSG272** | **34.2** | **ovale curtisi** | **37.0, 50.0** | **HC** | **Cameroon** | **Dschang** | **negative** | **96.9** | **95.3** | **94.4** | **67.4** |
| **SRR26037543poc7** | **N/A** | **ovale curtisi** | **N/A** | **sWGA** | **Cameroon** | **Imported to UK** | **negative** | **98.8** | **84.1** | **51.0** | **9.9** |
| **366152** | **29.4** | **ovale curtisi** | **34.8, -** | **HC** | **DRC** | **Bas-Uele** | **negative** | **97.1** | **95.3** | **94.3** | **131.0** |
| **364184** | **31.5** | **ovale curtisi** | **38.5, -** | **HC** | **DRC** | **Bas-Uele** | **negative** | **95.8** | **93.2** | **89.3** | **23.0** |
| **3073204** | **29.5** | **ovale curtisi** | **30.9, -** | **HC** | **DRC** | **Kinshasa** | **negative** | **96.1** | **94.3** | **93.1** | **76.2** |
| **1031006** | **32.3** | **ovale curtisi** | **35.3, -** | **HC** | **DRC** | **Kinshasa** | **positive** | **96.8** | **95.4** | **94.6** | **72.2** |
| **1013003** | **32.5** | **ovale curtisi** | **34.5, -** | **HC** | **DRC** | **Kinshasa** | **positive** | **96.2** | **94.5** | **93.2** | **62.5** |
| **1011201** | **34.0** | **ovale curtisi** | **35.1, -** | **HC** | **DRC** | **Kinshasa** | **positive** | **95.6** | **92.1** | **87.9** | **26.9** |
| **1012505** | **34.7** | **ovale curtisi** | **36.2, -** | **HC** | **DRC** | **Kinshasa** | **positive** | **95.5** | **92.5** | **89.3** | **30.8** |
| **2714 A** | **33.2** | **ovale curtisi** | **34.6, -** | **HC** | **Ethiopia** | **Amhara** | **negative** | **95.3** | **91.7** | **86.8** | **20.9** |
| **3116 A** | **31.8** | **ovale curtisi** | **32.5, -** | **HC** | **Ethiopia** | **Amhara** | **negative** | **95.9** | **94.0** | **92.4** | **34.4** |
| **ERR1739852pocgh01** | **N/A** | **ovale curtisi** | **N/A** | **LDB** | **Ghana** | **Upper East** | **positive** | **99.6** | **99.5** | **99.3** | **46.2** |
| **SRR26037545poc5** | **N/A** | **ovale curtisi** | **N/A** | **sWGA** | **Nigeria** | **Imported to UK** | **negative** | **99.4** | **94.8** | **73.5** | **13.6** |
| **SRR26037544poc6** | **N/A** | **ovale curtisi** | **N/A** | **sWGA** | **Nigeria** | **Imported to UK** | **negative** | **99.6** | **96.6** | **83.0** | **17.3** |
| **SRR26037542poc8** | **N/A** | **ovale curtisi** | **N/A** | **sWGA** | **Nigeria** | **Imported to UK** | **negative** | **99.5** | **98.5** | **92.3** | **22.2** |
| **ERR1428159povcu2** | **N/A** | **ovale curtisi** | **N/A** | **LDB** | **Nigeria** | **Imported to China** | **negative** | **99.6** | **99.3** | **98.6** | **62.2** |
| **SRR26037541poc9** | **N/A** | **ovale curtisi** | **N/A** | **sWGA** | **Sierra Leone** | **Imported to UK** | **negative** | **99.5** | **96.6** | **82.0** | **15.7** |
| **SRR26037546poc4** | **N/A** | **ovale curtisi** | **N/A** | **sWGA** | **South Sudan** | **Imported to UK** | **negative** | **99.6** | **99.4** | **99.2** | **85.8** |
| **1610 D0** | **32.2** | **ovale curtisi** | **34.8, -** | **LDB** | **Tanzania** | **Pwani** | **positive** | **99.4** | **94.9** | **71.2** | **14.6** |
| **426** | **34.0** | **ovale curtisi** | **40.0, -** | **LDB** | **Tanzania** | **Pwani** | **positive** | **99.5** | **99.2** | **98.6** | **82.3** |
| **475** | **35.6** | **ovale curtisi** | **41.1, -** | **LDB** | **Tanzania** | **Pwani** | **positive** | **97.6** | **75.8** | **41.3** | **10.4** |
| **1507081819_r2a** | **N/A** | **ovale wallikeri** | **N/A** | **sWGA** | **Cameroon** | **Imported to France** | **negative** | **95.8** | **78.0** | **53.9** | **16.3** |
| **1802062292_r4a** | **N/A** | **ovale wallikeri** | **N/A** | **sWGA** | **Cameroon** | **Imported to France** | **negative** | **98.5** | **97.7** | **95.0** | **63.9** |
| **ERR1739853powcr01** | **N/A** | **ovale wallikeri** | **N/A** | **LDB** | **Cameroon** | **Southwest** | **positive** | **98.4** | **96.1** | **75.7** | **12.2** |
| **SRR26037548pow16** | **N/A** | **ovale wallikeri** | **N/A** | **sWGA** | **Congo** | **Imported to UK** | **negative** | **98.5** | **96.5** | **86.0** | **19.0** |
| **111038** | **33.0** | **ovale wallikeri** | **-, 34.7** | **HC** | **DRC** | **Kinshasa** | **negative** | **97.0** | **94.4** | **88.5** | **425.6** |
| **113135** | **30.6** | **ovale wallikeri** | **44.4, 35.2** | **HC** | **DRC** | **Kinshasa** | **negative** | **98.5** | **98.2** | **97.9** | **547.8** |
| **353176** | **32.7** | **ovale wallikeri** | **45.4, 35.7** | **HC** | **DRC** | **Bas-Uele** | **negative** | **97.8** | **97.4** | **97.2** | **86.1** |
| **242012** | **32.9** | **ovale wallikeri** | **47.8, 40.7** | **HC** | **DRC** | **Sud-Kivu** | **negative** | **97.9** | **97.4** | **97.2** | **110.4** |
| **2090A** | **33.5** | **ovale wallikeri** | **-, 40.8** | **HC** | **Ethiopia** | **Amhara** | **negative** | **98.0** | **97.5** | **97.3** | **54.4** |
| **2562 DA** | **34.0** | **ovale wallikeri** | **-, 39.2** | **HC** | **Ethiopia** | **Amhara** | **negative** | **98.1** | **97.6** | **97.3** | **65.8** |
| **2589 DA** | **32.1** | **ovale wallikeri** | **-, 38.1** | **HC** | **Ethiopia** | **Amhara** | **negative** | **97.6** | **97.4** | **97.2** | **60.5** |
| **2680 A** | **31.5** | **ovale wallikeri** | **-, 38.3** | **HC** | **Ethiopia** | **Amhara** | **negative** | **98.2** | **97.8** | **97.6** | **142.6** |
| **2928 A** | **34.1** | **ovale wallikeri** | **-, 41.1** | **HC** | **Ethiopia** | **Amhara** | **negative** | **98.0** | **97.5** | **97.2** | **50.0** |
| **ERR1254543povwa2** | **N/A** | **ovale wallikeri** | **N/A** | **LDB** | **Gabon** | **Imported to China** | **negative** | **98.6** | **98.5** | **98.3** | **64.9** |
| **ERR1254542povwa1** | **N/A** | **ovale wallikeri** | **N/A** | **LDB** | **Gabon** | **Imported to China** | **negative** | **98.6** | **98.5** | **98.4** | **82.7** |
| **1503082061_r1a** | **N/A** | **ovale wallikeri** | **N/A** | **sWGA** | **Ivory Coast** | **Imported to France** | **negative** | **93.3** | **67.6** | **39.6** | **10.9** |
| **SRR26037551pow13** | **N/A** | **ovale wallikeri** | **N/A** | **sWGA** | **Kenya** | **Imported to UK** | **negative** | **98.5** | **94.9** | **82.2** | **20.7** |
| **SRR26037549pow15** | **N/A** | **ovale wallikeri** | **N/A** | **sWGA** | **Nigeria** | **Imported to UK** | **negative** | **98.6** | **98.3** | **97.8** | **71.6** |
| **1701100582_r3a** | **N/A** | **ovale wallikeri** | **N/A** | **sWGA** | **Senegal** | **Imported to France** | **negative** | **98.2** | **96.8** | **93.0** | **58.5** |
| **SRR26037550pow14** | **N/A** | **ovale wallikeri** | **N/A** | **sWGA** | **South Sudan** | **Imported to UK** | **negative** | **98.4** | **91.6** | **67.9** | **12.7** |
| **SOMO_ISA_041** | **30.0** | **ovale wallikeri** | **42.5, 38.4** | **HC** | **Tanzania** | **Songwe** | **negative** | **97.8** | **97.4** | **97.2** | **81.3** |
| **SOTU_TUN_019** | **29.6** | **ovale wallikeri** | **41.1, 37.2** | **HC** | **Tanzania** | **Songwe** | **negative** | **97.9** | **97.5** | **97.4** | **128.7** |
| **123** | **32.2** | **ovale wallikeri** | **-, 43.8** | **LDB** | **Tanzania** | **Pwani** | **positive** | **98.5** | **98.1** | **97.3** | **99.5** |
| **SRR26037552pow12** | **N/A** | **ovale wallikeri** | **N/A** | **sWGA** | **Tanzania** | **Imported to UK** | **negative** | **97.2** | **80.3** | **51.9** | **11.7** |

**Supplemental Table 1. Metadata of 21 *P. ovale curtisi* and 24 *P. ovale wallikeri* clinical isolates.** Technique refers to the method used to enrich *P. ovale* genomic DNA and/or remove human DNA. For Ct values, “-” indicates no amplification. *Pf* co-infection status was determined by real-time polymerase chain reaction. For 20 samples incorporated from other studies, *po18S* Ct values were not available. HC = hybrid capture; LDB = leukodepleted blood sample; sWGA = selective whole-genome amplification; DRC = Democratic Republic of the Congo; *Pf* = *P. falciparum;* _x cov. (%) = percent of species core genome covered by ≥_ reads (excluding *Poc* chromosome 10).

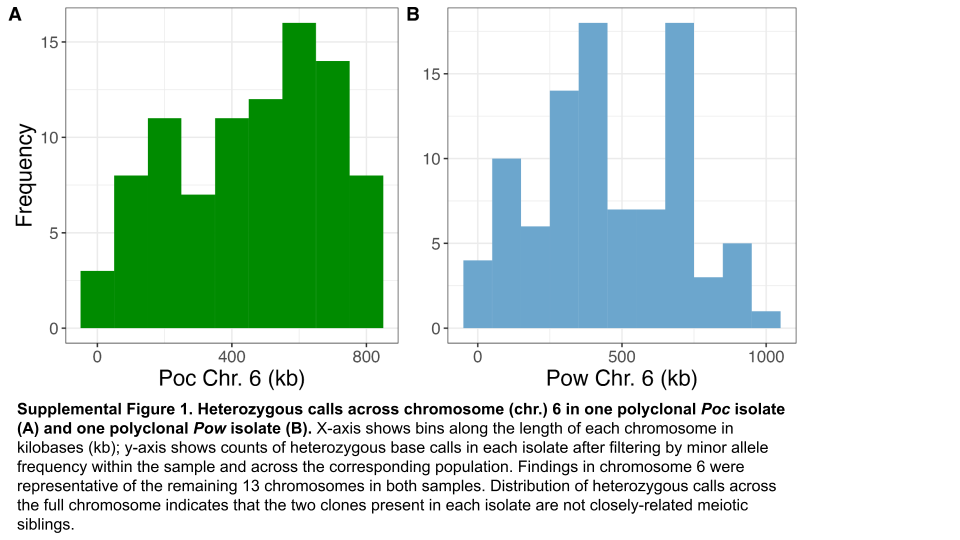

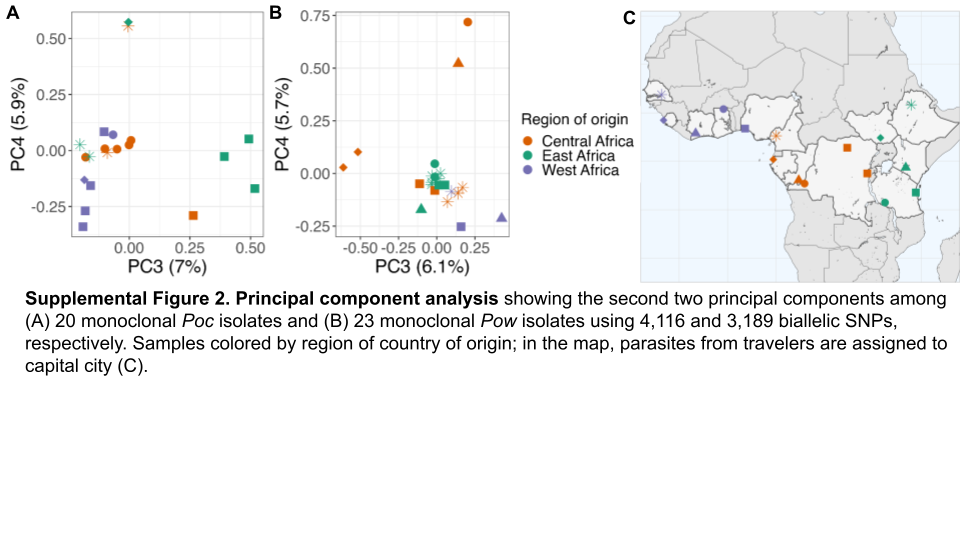

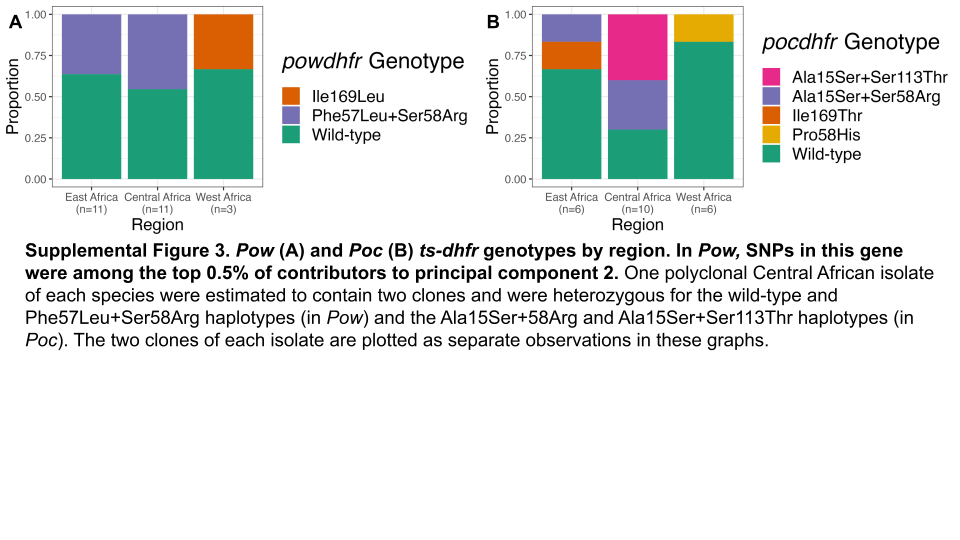

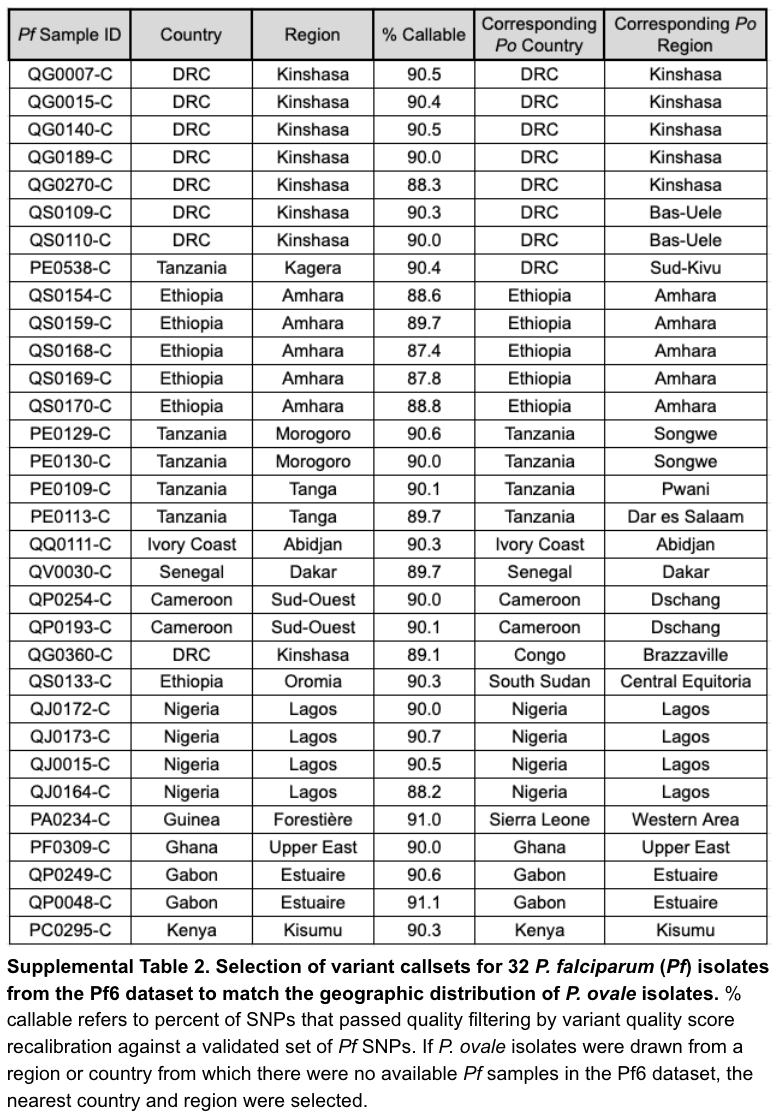

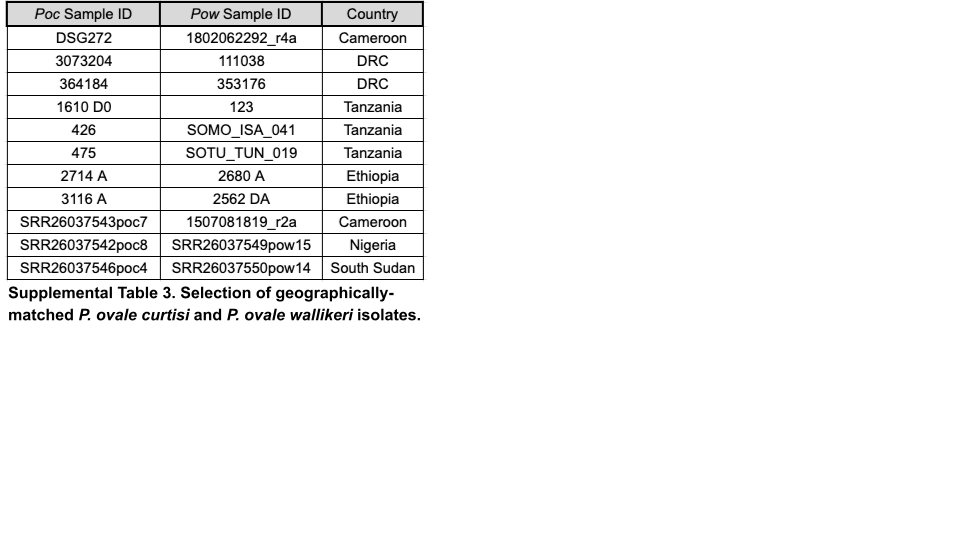

4.	Institute, B. ‘Picard Toolkit’, Broad institute, GitHub repository. *Picard Toolkit*.

5.	Van der Auwera, G. A. & O’Connor, B. D. *Genomics in the Cloud: Using Docker, GATK, and WDL in Terra*. (‘O’Reilly Media, Inc.’, 2020).

12.	Wickham, H. *ggplot2: Elegant Graphics for Data Analysis*. (Springer Science & Business Media, 2009).
